# Selection for greater dispersal in early life leads to faster age-dependent decline in locomotor activity and shorter lifespan

**DOI:** 10.1101/2022.07.03.498422

**Authors:** B G Ruchitha, Nishant Kumar, Chand Sura, Sudipta Tung

**Affiliations:** 1Integrated Genetics and Evolution Laboratory (IGEL), Department of Biology, Ashoka University, Sonipat, Haryana, India, 131029; 2Indian Institute of Science Education and Research (IISER) Pune, Pune, Maharashtra, India, 411008

## Abstract

Locomotor activity is one of the major traits that is affected by age. Greater locomotor activity is also known to evolve in the course of dispersal evolution. However, the impact of dispersal evolution on the functional senescence of locomotor activity is largely unknown. We addressed this knowledge gap using large outbred populations of *Drosophila melanogaster* selected for increased dispersal. We tracked locomotor activity of these flies at regular intervals until a late age. Longevity of these flies was also recorded. We found that locomotor activity declines with age in general. However interestingly, activity level of dispersal selected populations never drops below the ancestry-matched-controls, despite the rate of age-dependent decline in activity of the dispersal selected populations being greater than their respective controls. Dispersal selected population was also found to have shorter lifespan as compared to its control, a potential cost of elevated level of activity throughout their life. These results are crucial in the context of invasion biology as contemporary climate change, habitat degradation, and destruction provide congenial conditions for dispersal evolution. Such controlled and tractable studies investigating the ageing pattern of important functional traits are important in the field of biogerontology as well.

## Introduction

Dispersal is a central life-history trait (Bonte and Dahirel 2017). At an individual level, it can confer survival advantage against proximate environmental stresses (Studds et al. 2008), as well as reproductive advantage through a greater prospect of finding mates and suitable breeding grounds in the case of sexually reproducing organisms (Greenwood and Harvey 1982, Studds et al. 2008). At the population level, it modulates the rate and extent of adaptive dynamics by affecting the degree of gene flow between populations (Lenormand 2002, Garant et al. 2007). Dispersal also shapes local ecology by influencing processes such as territoriality, range expansion, species invasion, etc. (Hargreaves and Eckert 2014, Andrade-Restrepo et al. 2019).

It has been shown through both theoretical and empirical evidence that the dispersal abilities of individuals can evolve rather quickly (Perkins et al. 2013, Huang et al. 2015, Ochocki and Miller 2017, Tung et al. 2018a) in the presence of environmental conditions like climate change (Travis et al. 2013), habitat fragmentation (McPeek and Holt 1992, Berg et al. 2010), and habitat degradation and destruction (Fronhofer et al. 2014). These conditions are prevalent in the contemporary times due to, *inter alia*, climate change, human-induced landscape changes, and habitat destruction. Notably, dispersal is known to have strong associations with a number of traits of an individual, such as locomotor activity, aggression, body size, boldness, etc. (Wahlström 1994a, Michelangeli et al. 2017, Searcy et al. 2018, McGhee et al. 2021, Wu and Seebacher 2022), which are collectively known as dispersal syndrome (Clobert et al. 2009). Consequently, these traits also often evolve as a result of dispersal evolution (Fjerdingstad et al. 2007, Tung et al. 2018b). Such changes in organismal trait distributions can lead to the evolution of organisms with combinations of dispersal and dispersal syndromic traits with significantly higher potential for range expansion and biological invasion, and thereby threaten local ecological stability (Lambrinos 2004, Perkins et al. 2013, Renault et al. 2018). Thus, not surprisingly, investigating different aspects of dispersal evolution is a prime focus among contemporary ecologists, evolutionary biologists, and conservation biologists alike.

Despite the evident advantages that greater dispersal ability offers, such as avoiding local stressors, enhancing foraging opportunities, and facilitating mate finding, the diversity in dispersal properties among organisms in nature remains puzzling. This diversity can be understood by considering the costs associated with dispersal (reviewed in (Bonte et al. 2012)). These costs primarily arise from the energy-intensive nature of the dispersal process. Previous research involving pre-and post-dispersal analyses, has demonstrated that the dispersal process exacts a significant toll on an organism’s physiology (Mishra et al. 2022) and increases stress levels (Maag et al. 2019), although their long-term impacts across generations remain unclear. It follows logically that these physiological impacts would influence organismal performance later in life by accelerating the age-dependent decline in physiological functionality, also known as functional senescence of organismal traits (Grotewiel et al. 2005). Accelerated functional aging could diminish motor ability of an organism, thereby reducing its capacity to disperse and colonize new areas in later life, and may also alter age-dependent reproductive efforts and survival. For instance, functional senescence might result in decreased capacity for long-distance travel or increased vulnerability to predation during dispersal (Greenwood and Harvey 1982, Harvey et al. 1984, Newton 1993, Badyaev and Faust 1996, Forero et al. 1999, Pyle et al. 2001, Andreu and Barba 2006). These changes can significantly impact subsequent evolution of dispersal traits, thus providing a potential explanation for why we see diversity in dispersal properties among organisms in nature. Hence, it’s crucial to incorporate an understanding of senescence into studies on the evolution of dispersal, particularly concerning the functional senescence of traits associated with dispersal, or dispersal syndrome. Failure to do so could result in incomplete or inaccurate conclusions about the evolution of dispersal and its associated traits. However, the costs of dispersal evolution and its trade-offs with the functional senescence of dispersal syndromic traits are rarely investigated.

Locomotor activity is a prominent dispersal syndromic trait with strong association with greater dispersal ability across multiple species (Hanski et al. 2006; Cote et al. 2010; Fronhofer et al. 2015; Kosmala et al. 2017; Mishra et al. 2018). It is known to evolve as a correlated response to dispersal evolution (Tung et al. 2018b). However, the impact of this evolutionary adaptation on the functional senescence of locomotor activity remains unclear. Moreover, the relationship between age and locomotor activity varies among species. For example, some animals, such as senescence-accelerated mice (Miyamoto et al. 1986) and rabbits in open-field tests (Deyo et al. 1989) show an increase in activity with age; humans, mice, rhesus monkeys and some rats show a decrease in activity with age (Goodrick 1971, Dean et al. 1981, Emborg et al. 1998, McGibbon and Krebs 2004). In contrast, no significant change in activity with age has been seen in cats and some rats (Levine et al. 1987, Nagahara and Handa 1997). In the case of the insect model *Drosophila melanogaster*, (Le Bourg 1987) has shown an age-dependent decrease in activity in females, and an increase in activity in males till 5 weeks of age, after which activity declines. However, some studies have shown that such sexual dimorphism in the age-dependent activity of *Drosophila* is strongly influenced by the genotype of the flies used (van Dijken et al. 1979, Fernández et al. 1999). Therefore, it is not yet clear how mobility changes with age in sexually dimorphic organisms, such as fruit flies *Drosophila melanogaster*. Given that males and females of sexually dimorphic organisms tend to defer with respect to their resource allocation strategies in reproduction and somatic maintenance as well as differ in metabolic rates, it is likely that selection for dispersal ability can shape functional senescence in males and females differently (Garratt 2020). However, there is limited understanding of how evolution for increased dispersal ability affects different sexes at various ages.

In this study, we aimed to fill this knowledge gap by investigating locomotor activity and lifespan of male and female individuals from four large, outbred populations selected for greater dispersal ability and propensity (Tung et al. 2018a). We examined the locomotor activity of the populations selected for greater dispersal and their ancestry-matched controls after maintaining both types of populations under identical conditions for nine generations, or approximately five months, without any further selection for dispersal. The age-dependent locomotor activity patterns of the evolved and control populations are compared for males and females at multiple time points across the lifespan to understand the impact of dispersal evolution on the functional senescence of locomotor activity in these flies. We began by analysing the aging profile in a cross-sectional study to understand how the trait has changed in these populations. Next, considering variation in lifespan among individuals within a population and to eliminate potential survivorship bias because of it, we conducted a longitudinal study by following flies individually and measuring their locomotor activity across age. Additionally, in this longitudinal study, we compared the longevity of the evolved and control populations to understand the impact of functional aging of locomotor activity on demographic aging by measuring lifespan.

Through these experiments, we provide insights on functional senescence of locomotor activity and demographic senescence in the two sexes and how the evolution of dispersal affects them. These aspects are particularly relevant as locomotor activity plays a critical role in reproduction, encompassing activities such as mate searching and successful courtship for males, as well as locating suitable egg-laying surfaces for females, ultimately influencing their post-dispersal colonization success.

## Methods

### Experimental Populations

We used four large (n = ∼2400), outbred, laboratory-maintained populations of *Drosophila melanogaster* (VB_1-4_) that were selected for greater dispersal propensity and ability for 153 generations (>6 years), and their respective ancestry-matched control (VBC_1-4_) populations. For clarity and ease of reading, henceforth, we will refer to the VB populations as ‘dispersal-selected’, ‘selected’, or ‘D’ populations interchangeably. Similarly, the VBC populations will be referred to as ‘control’ or ‘C’ populations. These populations were derived from a common ancestry, therefore had comparable genetic architecture, and were maintained identically under standard laboratory conditions. Details of the selection protocol and maintenance regime of these populations were reported previously (Tung et al. 2018a). Briefly, the D_1-4_ and C_1-4_ populations were maintained in 15-day discrete generation cycles on a standard banana-jaggery medium (following the recipe of Sheeba and Joshi 1998) at 25°C, with constant light. On the 12^th^ day from the day of egg collection, only the most dispersive half of the population were artificially selected. This was done by subjecting the D_1-4_ populations to a three-compartment source-path-destination dispersal selection setup, where the flies from D_1-4_ populations were placed in the source compartment, allowed to disperse via the path to reach the destination compartment. Among these flies, only the first ∼50% of the population estimated visually to reach the destination compartment were allowed to lay eggs for the next generation. To maintain the consistent breeding population size of the D_1-4_ populations across generations while also not biasing the density of the population in the source compartment, two such source-path-destination setups would be created for each population of the D_1-4_ populations. Thus, collecting approximately the first 50% of the population from each of the two source-path-destination setups ensured a consistent breeding population size in the selected populations comparable to that of the corresponding control populations. These populations were maintained at a high population size of approximately 2400 flies per population to minimize the impact of inbreeding. C_1-4_ populations were not subjected to selection for dispersal, but the rest of the rearing conditions were identical to that of the D_1-4_ populations. To obtain the next generation of flies, in both D and C populations, eggs were collected on the 15^th^ day post egg collection, ensuring that no differential age-dependent selection is acting on the D and C populations.

## Experiments

### Experiment 1: For investigating population level effect

Prior to this experiment, selection pressure for dispersal was relaxed for eight generations i.e. D_1-4_ and C_1-4_ populations were maintained identically without selection pressure to minimize non-genetic parental effects. ∼300 eggs were collected in drosophila-bottles (Laxbro® #FLBT20) containing 50mL banana-jaggery food. Eight such replicates were made for each population. On 11^th^ day from egg collection, all flies from eight bottles of same population were pooled into a single cage (25 cm × 20 cm × 15 cm) and maintained with fresh food being provided every alternate day. This was done for each of the eight populations (D_1-4_ and C_1-4_). From each of these population cages, 32 flies of each sex were randomly sampled at predetermined age-points (see “Assay Details” below). Overall, locomotor activity of 2048 flies (32 flies × 2 sexes × 8 populations × 4 age-points) was measured for this experiment.

Figure S1 shows a schematic diagram of the experimental design for investigating population level effect.

### Experiment 2: For tracking individual flies

In experiment 2, we collected longitudinal data as opposed to the cross-sectional data gathered in experiment 1. Since the lifespans of individuals can vary within a population, cross-sectional activity data can be affected by survivorship bias, which can influence the interpretation of results. Therefore, experiment 2 was designed as a proof-of-principle validation of the findings from experiment 1, and due to logistical constraints was conducted using one set of ancestry-matched selected and control populations (D_2_ and C_2_), chosen at random.

This experiment was performed on the dispersal selected population, after relaxation of selection pressure for dispersal over nine generations, and its ancestry matched control. In order to generate experimental flies for each population, ∼300 eggs were collected in drosophila-bottles (Laxbro® FLBT20) containing 50mL banana-jaggery food. Three such replicates were made for each population. From the three bottles, 30 individuals of each sex were chosen at random during the P13-P15 stage of pupa and hosted individually in glass vials containing ∼4mL of food and incubated at 25°C. On eclosion, these 120 virgin adult flies (*i.e.* 30 individuals per sex per population) were checked daily for survival until death, and their age-dependent locomotor activity was tracked by measuring the trait at predetermined age-points (see “Assay Details” below). The flies were provided fresh food every alternate day, while strictly maintaining their vial identity. Figure S2 shows a schematic diagram of the experimental design for tracking individual flies.

## Assay Details

### Locomotor activity assay (used for experiments 1 and 2)

Locomotor activity was measured as the number of times per hour that each fly present in a glass tube (diameter: 5 mm, length: 8 cm) crossed the middle of the tube. This was done using Drosophila Activity Monitoring (DAM2) data collection system (Trikinetics Inc, Waltham, MA) which recorded the number of times each fly would cross the infrared beams at the centre of the glass tube every 30 seconds using a standard protocol (Chiu et al. 2010).

To minimize any potential bias from variation in movement patterns of the flies, all treatments in our experiment were handled consistently.

For experiment 1, we tracked locomotor activity of four independent but identically maintained, large, outbred, dispersal-selected populations D_1-4_ across age and contrasted the same against corresponding four ancestry-matched control populations C_1-4_. To do this, we randomly sampled 32 flies of each sex from each of these eight populations (∼2400 adults on the 12^th^ day in each population) on the 12^th^, 26^th^, 40^th^, and 47^th^ day from the day of egg collection and measured their locomotor activity. On the specified days of activity measurement, flies were collected from population cages and introduced into clean glass tubes (diameter: 5 mm, length: 8 cm), without CO_2_ anaesthesia. Each glass tube was secured with cotton plugs on both sides. Activity was recorded for 2h and 5mins, and the initial 5 min was excluded to minimize the effect of disturbance caused at the time of setup. At the end of the recording, mortality was recorded and all the flies in the glass tubes were discarded. Overall, for experiment 1, locomotor activity of 256 flies (32 flies × 2 sexes × 4 age-points) of each population was measured.

For experiment 2, we repeatedly measured locomotor activity of same set of 30 male and 30 female flies each of dispersal-selected and control populations on the 12^th^, 26^th^, 33^rd^, 40^th^, and 47^th^ day from the day of egg collection. On the specified days of activity measurement, flies were introduced into individually labelled (to maintain identity of each fly), clean glass tubes (diameter: 5 mm, length: 8 cm), without CO_2_ anaesthesia. Each glass tube contained 1.5% w/vol agar, and was sealed with parafilm on one end and secured with cotton plug on the other. Activity was recorded for 4h and 5mins, and the initial 5 min was excluded to minimize the effect of disturbance caused at the time of setup. At the end of the recording, the flies were transferred back to the corresponding individually labelled glass vials (to maintain identity of each fly) with food. Overall, for experiment 2, 60 flies (30 flies × 2 sexes) of each population were individually tracked by maintaining them in labelled containers and their locomotor activity was measured at five pre-determined age-points.

### Longevity assay (used for experiment 2)

Longevity of each fly was assessed as the number of days it was alive. Each day, we manually looked at each of the flies that were being individually tracked (from experiment 2) until all flies were dead. Complete absence of movement of limbs, abdomen and proboscis was recorded as death.

## Statistical Analysis

### Locomotor activity assay of experiment 1

We disregarded the data of the flies which were found dead at the end of the recording (details of sample size is present in Table S1). For all the remaining flies, out of the 2 hours of recording, we only considered the first 1 hour of data. We did this to minimize the effect of potential artifacts generated by inactivity due to death and hyperactivity due to starvation (Yu et al. 2016). From the data obtained after these corrections, we calculated the average of one hour locomotor activity of 32 flies for each of 16 treatment groups (2 sexes × 8 populations) across 4 age-points.

To test how evolution of greater dispersal affects age-dependent locomotor activity of dispersal selected population, we analysed the locomotor activity data using a linear mixed effects model with Selection (dispersal-selected vs. control), Sex (male vs. female), Age (12, 26, 40, and 47) as fixed factors, and population identity (1, 2, 3 and 4) as random factor. We used ‘lmer’ function from ‘lme4’ package in R v4.3.2 with a model *Activity ∼ Selection+ Sex+ Age + (1|Block) + Selection*Sex + Selection*Age + Sex*Age*. When the main or interaction effect was significant, post-hoc was performed using Tukey’s HSD test using ‘emmeans’ function from ‘lsmeans’ package in R v4.3.2.

### Locomotor activity assay of experiment 2

The 4h locomotor activity data was used to calculate per hour activity. In order to distinguish between within-individual variation in locomotor activity across age and between-individual variation in locomotor activity resulting from differences in longevity, we divided age into two components: Delta age and Average age, respectively (van de Pol and Wright 2009).

Here, Average age represents the mean of all timepoints for an individual, while Delta age reflects timepoint specific deviations from the average. Now, to test how evolution of greater dispersal affects locomotor activity of individual flies (*i.e.* here, we measure longitudinal data as opposed to the cross-sectional data obtained from experiment 1) with age, the per hour activity data was subjected to a linear mixed effects model with Selection (dispersal-selected vs. control), Sex (male vs. female), Average age and Delta age as fixed factors, and identity of individual flies as random factor. If a significant Selection × Delta age interaction is observed, it would imply that the selection for increased dispersal is associated with the selection for accelerated locomotor senescence.

### Longevity assay of experiment 2

To check whether there is change in lifespan due to evolution of greater dispersal, we performed a full-factorial 2-way fixed factor ANOVA on longevity data (measured as mention in “Assay Details” section above) with Selection (dispersal-selected vs. control) and Sex (male vs. female) as fixed factors using STATISTICA® v5 (StatSoft. Inc., Tulsa, Oklahoma) software.

Additionally, we conducted a comparison of survivorship curves employing a Cox proportional-hazards model. The ‘Surv’ function from the ‘survival’ package in R v4.3.2 was used to construct the survival object, incorporating both time and event status. A full factorial model, incorporating Selection (dispersal-selected vs. control) and Sex (male vs. female) as fixed factors, along with their interaction was fitted using the ‘coxph’ function from the same R package, and summary statistics were extracted. The proportional hazards assumption in the Cox proportional hazards model was assessed graphically by checking the parallelism of the log-minus-log survival plots created using the ‘survfit’ and ‘plot’ functions in R. The relatively parallel lines suggest that the proportional hazards assumption is reasonably met (Figure S3).

To discern whether dispersal-selection impacts longevity through baseline mortality or age-dependent mortality rate, we fitted a two-parameter Gompertz function using the ‘flexsurvreg’ function from the ‘flexsurv’ package in R v4.3.2. We conducted separate analyses for males and females in both the selected and control populations. In this analysis, the shape and rate parameters correspondingly signify the baseline mortality early in life and the age-dependent mortality rate for each group of individuals.

To determine if there is a correlation between locomotor activity and lifespan within treatment groups, fitted a linear model for lifespan with Average locomotor activity, Selection, Sex and their interaction as fixed effects (*Longevity ∼ Selection*Sex*AvgActivity*) using ‘lm’ function from R v4.3.2. The same analysis was also repeated with locomotor activity measured at the first time point as a fixed factor instead of Average locomotor activity of the individuals.

All the graphs were generated using *matplotlib* (Version: 3.5.1) module in Python 3.8.3 (https://www.python.org/).

### Dispersal evolution leads to greater rate of functional senescence

In the cross-sectional study (experiment 1) of locomotor activity across age, although activity declines in both dispersal selected populations and their ancestry-matched controls, we found a significant interaction between Age and Selection (Figure 1A; Age × Selection *p* = 2 ×10^-22^) with faster decline in locomotor activity in the dispersal-selected populations than in the ancestral control populations. This observation is further validated, when we tracked locomotor activity of individual flies from both dispersal-selected and control populations across age in an independent longitudinal study i.e., experiment 2 (Figure 2A). For the analysis of this data, we split age into a Delta age and an Average age component to delineate the effect of within-individual variation in locomotor activity across age and between-individual variation in locomotor activity due to difference in longevity, respectively. We found a significant Selection × Delta age interaction (Figure S5, *p* = 2×10^-11^) demonstrating that selection for greater dispersal is accompanied with selection for accelerated locomotor senescence. Further, we found that the main effect of Average age (*p* = 0.4) and Selection × Average age interaction (Figure S6, *p* = 0.68) are not significant, indicating that the above observation is not due to any variation in longevity between the individuals or survivorship bias in the experiment. This is further validated as we found that there is no significant correlation between average locomotor activity and longevity of the individuals (F_1, 103_ = 0.07, *p* = 0.79) highlighting that increased locomotor activity is not linked with reduced lifespan within treatment groups. The result remained same when we evaluated correlation between early-life activity with longevity of the individuals (F_1, 103_ = 0.01, *p* = 0.91).

**Figure 1.**
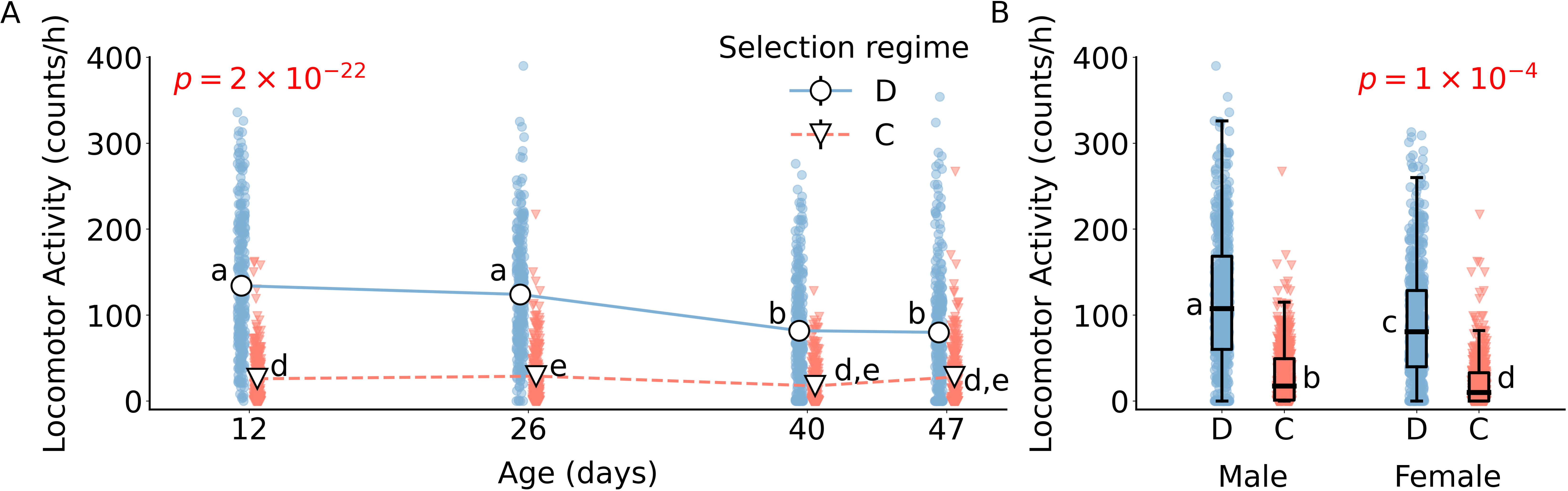
**Age-and sex-dependent locomotor activity profile from cross-sectional data.** (A) Mean (± 95% CI) locomotor activity of populations from dispersal-selected D (circle) and its ancestry-matched control C (triangle) populations are plotted against age, measured in days from the time of egg collection. Activity level of each population at the tested age-point of D and C populations are represented as blue and red scatter points respectively. Some of the error bars are not visible due to their small size. *p* value for Selection × Age interaction is mentioned in red font. (B) Box-plot representing locomotor activity of dispersal-selected D and its ancestry-matched control populations across males and females. The edges of the box denote 25th and 75th percentiles, while the black solid line represents the median. The blue and red scatter points represent the data for all the replicates of D and C populations respectively. *p* value for Selection × Sex interaction is mentioned in red font. Different lower-case alphabets denote statistically significant differences (p<0.05).

**Figure 2.**
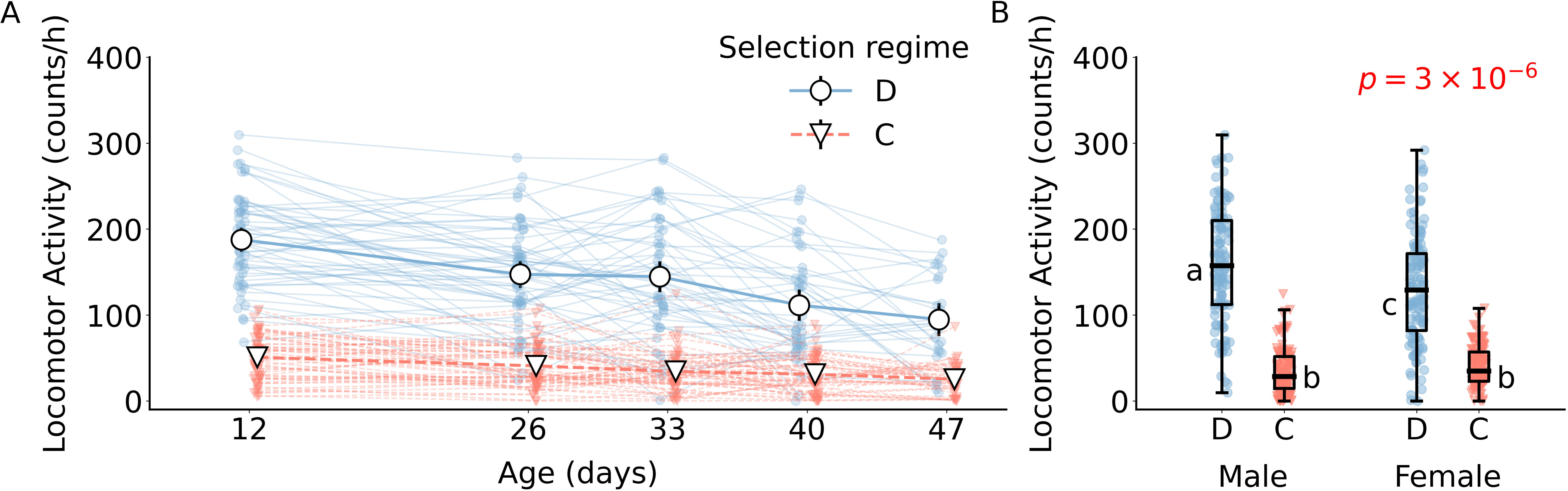
**Age-and sex-dependent locomotor activity profile from longitudinal data.** (A) Mean (± 95% CI) locomotor activity of individuals from dispersal selected population D (circle) and its ancestry-matched control population C (triangle) are plotted against age, measured in days from the time of egg collection. Activity level of each individual at the tested age-point of dispersal selected and control populations are represented as red and blue scatter points respectively. Scatter points for same individual are connected by solid line. (B) Box-plot representing locomotor activity of individuals from dispersal-selected D and its ancestry-matched control C populations across males and females. The blue and red scatter points represent the data for all the replicates of D and C populations respectively. The edges of the box denote 25th and 75th percentiles, while the black solid line represents the median. p value for Selection × Sex interaction is mentioned in red font. Different lower-case alphabets denote statistically significant differences (p<0.05). Some of the error bars are not visible due to their small size.

Interestingly, our cross-sectional analysis showed a significant Age × Sex interaction (Figure S4, p = 5 × 10^-8^). Additionally, our longitudinal investigation revealed a significant Sex × Average age interaction (*p* = 0.02), whereas the Sex × Delta age interaction (*p* = 0.33) was not significant. This suggests that the observed sex-specific variation in locomotor activity across age points predominantly stem from variations in lifespan between sexes rather than differences in the rate of senescence of locomotor activity.

### Despite elevated functional senescence, dispersal selected flies have high locomotor activity across age

We see that despite dispersal evolution causing greater rate of decline in locomotor activity with age (as mentioned above), the locomotor activity of the dispersal selected flies does not drop below the activity level of their ancestry-matched control flies even at late age in both cross-sectional (experiment 1, Figure 1A and Table S1) and longitudinal (experiment 2, Figure 2A and Table S2) studies. This observation is consistent across both males and females. However, interestingly, the increment in locomotor activity in selected males was greater than that in females, leading to a significant Selection × Sex interaction in both the experiment 1 (Figure 1B, *p* = 1 × 10^-4^) and experiment 2 (Figure 2B, *p* = 3 × 10^-6^). A comprehensive table containing mean activity levels and *p* values for experiment 1 and experiment 2 can be found in Table S1 and Table S2.

### Selection for greater dispersal properties trades-off with lifespan

In experiment 2, as we tracked each fly individually, we were able to measure the lifespan of these flies. From this data, we see that the dispersal selected flies live significantly shorter (Figure 3 and Table S3; F_1,107_ = 4.52, p = 0.036) than their ancestry-matched control. We also see that longevity between the dispersal-selected and ancestral population does not vary differentially due to sex i.e. the difference in longevity between the dispersal selected and their ancestry-matched control flies is comparable in both males and females (Selection × Sex F_1,107_ = 0.76, p = 0.4). We arrived at the same conclusion when we compared the survivorship curves through a cox proportional hazards model, yielding a significant main effect of Selection (Figure 3A, *p* = 5 × 10^-4^) and a non-significant effect of Selection × Sex interaction (*p* = 0.23). Interestingly, fitting the lifespan data of the selected (D) and control (C) populations with a two-parameter Gompertz function revealed that the reduced longevity in the selected populations primarily results from an accelerated age-dependent mortality rate.

**Figure 3.**
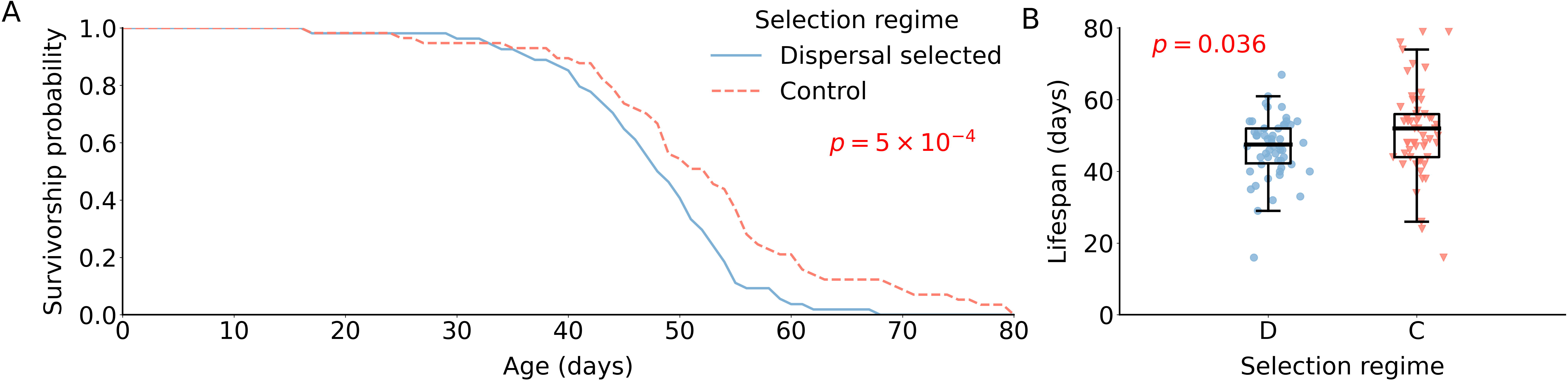
**Survivorship curve and lifespan of dispersal-selected and control populations.** (A) Survivorship probability (i.e. proportion of viable individuals) of dispersal-selected D (blue solid line) and control C (red dashed line) populations across time in days. p value for the main effect of Selection as per the Cox proportional hazard model is mentioned in red font. (B) In the box plot, the middle line, lower and upper edge of the box, lower and upper end of the whiskers represent mean, 25th, 75th, 5th, 95th percentile of the longevity data respectively. The scatter points denote longevity of each individual of the dispersal-selected D (blue circles) and control C (red triangles) populations. ANOVA p value for the main effect of Selection is mentioned in red font.

This result is consistent in both males (shape parameter: 0.14 for selected vs. 0.07 for control) and females (shape parameter: 0.18 for selected vs. 0.097 for control). In contrast, the baseline mortality in early life was actually lower in the selected population, as indicated by the rate parameter values: -9.322 for selected males compared to -6.853 for control males, and -10.037 for selected females compared to -7.515 for control females.

## Discussion

In this study, we have used four large (n ∼2400) outbred populations of *Drosophila melanogaster*, which were selected for greater dispersal at their early life. In these dispersal-evolved populations, we quantified the status of functional senescence in locomotor activity of both male and female flies, and contrasted the data against their ancestry matched control populations, which haven’t undergone dispersal evolution. Interestingly, we found that dispersal evolution leads to faster decline in locomotor activity with age (Figure 1) indicating that evolving greater dispersal at early life, indeed, has adverse effect in the late life stages resulting in accelerated functional senescence. Further, in order to demonstrate that the above result is not an artefact of cross-sectional experiment involving population level averages, as a proof-of-principle, we tracked individual flies throughout their lifespan and measured locomotor activity at predetermined age-points. This experiment supported the above insight by yielding the same results (Figure 2).

Moreover, we also found that despite faster age-dependent decline in locomotor activity in dispersal evolved flies, their locomotor activity does not drop below the activity level of their ancestry-matched control flies at any given age (Figure 1A and Figure 2A). It is further noteworthy that this elevated locomotor activity in the dispersal selected flies persisted even in the absence dispersal selection for over nine generations. Together, these imply that the dispersal evolved individuals continue to carry and exhibit correlated traits of dispersal evolution like increased locomotor activity even after they reach their destination, and it can last over multiple generations long after the selection pressure for dispersal ceases. At the mechanistic level, these observations indicate that the metabolic remodelling as a correlated response to dispersal evolution proposed in previous study (Tung et al. 2018b) is likely to have become constitutive in the dispersal selected flies leading to persistent elevated expression of locomotor activity in the selected flies throughout their life. Elevated locomotor activity in the more dispersive individuals throughout their life have important ecological implications as well. Previous studies have indicated that increased locomotor activity enhances mating success by elevating courtship behaviour (Kyriacou 1981, Cobb et al. 1987) and assists females in locating suitable oviposition sites (Ferguson et al. 2015). Consequently, under similar circumstances, the heightened locomotor activity observed in dispersal-selected populations would theoretically yield greater reproductive success in a natural environment, thus contributing to greater evolutionary success. However, we also note a shorter lifespan in dispersal-selected populations, which could potentially counterbalance the reproductive advantages conferred by increased locomotor activity. Consequently, determining the net impact of dispersal evolution on overall reproductive output across the lifespan is not straightforward. Additionally, it is important to keep in mind that the sustained elevated locomotor activity in the dispersal-selected populations we see in our study are from laboratory conditions of *ad libitum* food and absence of predation. The environmental conditions in the wild would be much harsh and thus sustained elevated locomotor activity may not be selected for in wild populations. Therefore, further studies are required to understand the nuanced effect of dispersal selection on reproductive and evolutionary success of populations selected for greater dispersal.

Furthermore, the observed faster functional senescence and shorter lifespan in dispersal-selected flies could potentially be a negative consequence of constitutively elevated expression of locomotor activity in the selected flies throughout their life (Figure 1A and Figure 2A). Although we do not have direct evidence for a mechanistic explanation behind these observations, the following mechanisms might be responsible for this. Firstly, the dispersal-selected flies were previously reported to have evolved an elevated level of metabolite 3-HK (Tung et al. 2018b), which is associated with age-dependent neural degeneration in *Drosophila* (Savvateeva et al. 2000) and hence the greater rate of neuromuscular impairment, leading to deterioration of locomotor activity and overall reduced lifespan. Secondly, studies have shown that elevated physical activity can lead to the activation of stress-responsive pathways such as the MAPK and NF-_κ_B pathways (Kramer and Goodyear 2007, Egan and Zierath 2013), which can lead to increased oxidative stress and inflammation. Similarly, the selected flies were found to have a greater level of cellular respiration resulting in an elevated level of ATP in these flies (Tung et al. 2018b). It is also known that free radicals are produced during ATP production via oxidative phosphorylation in mitochondria (Cadenas and Davies 2000). These responses can ultimately lead to the activation of pro-aging pathways and the observed acceleration of aging-associated declines in physiological function in the selected populations. Thirdly, *Drosophila* insulin-TOR signalling pathways are known to affect cardiac functional aging, and thereby shorten lifespan (Wessells et al. 2009). Systemic glucose level is reported to be higher in dispersal-selected flies (Tung et al. 2018b), which can lead to relatively greater activation of insulin-TOR signalling pathways in these flies resulting in faster senescence.

While further targeted experiments are necessary to fully understand the exact underlying mechanisms, this study presents the first empirical evidence that aging evolves as a direct consequence of dispersal evolution. Moreover, the metabolic changes mentioned above emerged in response to elevated cellular respiration to meet the increased energy demands for dispersal in early life (Tung et al., 2018b). The evolutionary theories of aging suggest that aging evolves as a result of weakened selection pressure later in life (Medawar 1946; Williams 1957). Given that, for the artificial selection procedure, eggs from these flies are collected within three days after dispersal, the later part of life remain under selection shadow. Thus, it is conceivable that over the course of evolution, the evolved populations have accumulated alleles that facilitate greater dispersal in early life, albeit at the expense of negative pleiotropic effects in late life, and increasing the likelihood of death. Thus, this study serves as an empirical demonstration of antagonistic pleiotropy, a population-genetic concept to explain the evolution of aging based on the broader idea of evolutionary constraints and life history tradeoffs (Medawar 1946; Williams 1957). Even in natural population of neotropical butterflies, higher dispersal rates led to decreased longevity (Tufto et al. 2012). On the other hand, from the existing literature, we also know that often dispersal, particularly in severely patchy habitats, results in populations with small effective population size, which leads to strong random genetic drift in these population (Polechová 2022), which can result in reduced lifespan and accelerated aging, as seen in a study with water fleas, *Daphnia magna* (Lohr et al. 2014). However, in a natural setting, it is difficult to dissect out whether reduced lifespan and accelerated senescence is due to dispersal or due to genetic drift caused by habitat fragmentation and/or isolation. Whereas in our study, using large outbred populations we could show that, independent of genetic drift route, dispersal evolution can have direct negative impact on the physiological function of organisms at late life and their lifespan.

Moreover, it is worth noting that in a previous study on the same populations (Tung et al. 2018a), the difference between the selected and control populations in terms of dispersal ability and propensity was apparent as early as after 10 generations of selection. However, the lifespan of the dispersal-selected and control populations was not significantly different, although there was a trend (Figure 3C in Tung et al. 2018b). Whereas, the same trade-off became apparent after 153 generations of dispersal selection. This demonstrates the fact that the rate of evolution of the correlated traits can differ drastically compared to the traits which are under direct selection. Further, it also highlights that the trait correlations observed in the short-term can change over a longer evolutionary timescale.

Finally, our findings demonstrate that although the trade-off between dispersal and longevity logically indicates a rapid decline of dispersal and associated traits such as locomotor activity upon relaxation of selection for greater dispersal, it may not occur as per the expectations in all circumstances. Specifically, we observed elevated levels of locomotor activity in our dispersal-evolved flies even after nine generations of relaxation of selection pressure. This apparent discrepancy can be explained by the fact that these flies are maintained in a 15-day discrete-generation lifecycle, and therefore, they do not experience their full lifespan in these laboratory conditions. Thus, one may speculate that the trade-off may be avoided in natural settings where organisms reproduce rapidly at an early stage of life. This hypothesis can be tested in future research using natural populations. Furthermore, the effect of faster aging and shortened lifespan on other life-history and physiological traits is an important avenue for future research to explore.

In summary, evolutionary history has an impact on age-dependent locomotor activity in fruit flies. The flies that evolved greater dispersal at their early life suffer from steeper age-dependent decline in locomotor activity, and have shorter lifespan. This pattern of trade-off between dispersal and senescence is observed in both male and females. The exact molecular mechanism behind this antagonistic effect (shorter lifespan as a result of selection for dispersal in early life) remains to be discovered. Additionally, we found that a correlated increase in locomotor activity due to dispersal selection does not disappear even after nine generations of removal of selection pressure and the increased locomotor activity remains higher than that of the control populations until later ages. Please note that locomotor activity is not the same as dispersal. In our experiment here, locomotor activity change is a correlated response to dispersal selection. In the current global scenario, where habitat degradation and destruction, and rapid climate change are rampant, species that are better at dispersing have a distinct evolutionary advantage. This highlights the importance of gaining a complete understanding of how dispersal evolution and senescence influence each other. As senescence greatly affects the performance of an organism, and thereby impacts its long-term persistence and adaptation, such knowledge can inform us about the distribution and interactions of species with their environment, and have far-reaching implications for various eco-evolutionary processes. For example, it can help us predict the success of species in colonizing new areas, invading new territories, expanding their range, and spreading diseases, as well as the stability of populations with strong dispersal abilities. Thus, this understanding is pertinent and relevant to the research community, especially those working in the fields of evolution, ecology, and conservation.

## Acknowledgements

The authors thank Prof. Sutirth Dey from Indian Institute of Science Education and Research (IISER)-Pune for kindly providing us the experimental populations. The authors also thank Prof. Dey and Akhila Mudunuri for their valuable comments on an earlier version of the manuscript. RBG acknowledges the support from Kishore Vaigyanik Protsahan Yojana (KVPY) fellowship, NK and CS acknowledge the support of the Research and Development Office, Ashoka University. ST acknowledges the support of DBT/Wellcome Trust India Alliance Early Career Fellowship (#IA/E/18/1/504347) and Ashoka University.

## Conflict of interest

The authors declare no conflict of interest.

## Notes

### Competing Interest Statement

The authors have declared no competing interest.

### Summary of Updates

Analysis, fugures and the main text are updated.

