## Supplementary for "Selection for greater dispersal in early life leads to faster age-dependent decline in locomotor activity and shorter lifespan"

for

**Table S1. Descriptive statistics of locomotor activity for VB and VBC populations at all age points in Experiment 1**

| Selection | Sex | Age | Block | Average Activity (counts/h) | Sample size |
| --- | --- | --- | --- | --- | --- |
| VB | M | 12 | 1 | 192.5 | 32 |
| VB | M | 26 | 1 | 163.75 | 32 |
| VB | M | 40 | 1 | 148.71875 | 32 |
| VB | M | 47 | 1 | 178.25 | 31 |
| VB | F | 12 | 1 | 196.4375 | 32 |
| VB | F | 26 | 1 | 129.125 | 32 |
| VB | F | 40 | 1 | 71.3125 | 32 |
| VB | F | 47 | 1 | 66.8125 | 30 |
| VBC | M | 12 | 1 | 43.28125 | 31 |
| VBC | M | 26 | 1 | 38.4375 | 32 |
| VBC | M | 40 | 1 | 27.3125 | 30 |
| VBC | M | 47 | 1 | 29.15625 | 25 |
| VBC | F | 12 | 1 | 42.40625 | 32 |
| VBC | F | 26 | 1 | 30.875 | 32 |
| VBC | F | 40 | 1 | 10.46875 | 32 |
| VBC | F | 47 | 1 | 20.75 | 29 |
| VB | M | 12 | 2 | 195.0625 | 32 |
| VB | M | 26 | 2 | 173.90625 | 32 |
| VB | M | 40 | 2 | 119.875 | 31 |
| VB | M | 47 | 2 | 81.53125 | 24 |
| VB | F | 12 | 2 | 183.5 | 32 |
| VB | F | 26 | 2 | 161.84375 | 32 |
| VB | F | 40 | 2 | 76.28125 | 31 |
| VB | F | 47 | 2 | 78.4375 | 32 |
| VBC | M | 12 | 2 | 21.40625 | 32 |
| VBC | M | 26 | 2 | 40.09375 | 32 |
| VBC | M | 40 | 2 | 30.40625 | 29 |
| VBC | M | 47 | 2 | 55.34375 | 28 |
| VBC | F | 12 | 2 | 25.28125 | 32 |
| VBC | F | 26 | 2 | 20.78125 | 31 |
| VBC | F | 40 | 2 | 14.125 | 32 |
| VBC | F | 47 | 2 | 17.875 | 22 |
| VB | M | 12 | 3 | 59.875 | 32 |
| VB | M | 26 | 3 | 110.84375 | 32 |
| VB | M | 40 | 3 | 65.40625 | 32 |
| VB | M | 47 | 3 | 77.8125 | 19 |
| VB | F | 12 | 3 | 75.84375 | 32 |
| VB | F | 26 | 3 | 90.28125 | 32 |
| VB | F | 40 | 3 | 50.125 | 32 |
| VB | F | 47 | 3 | 33.6875 | 22 |
| VBC | M | 12 | 3 | 19.6875 | 32 |
| VBC | M | 26 | 3 | 23.15625 | 32 |
| VBC | M | 40 | 3 | 19.4375 | 29 |
| VBC | M | 47 | 3 | 28.5 | 20 |
| VBC | F | 12 | 3 | 11.625 | 32 |
| VBC | F | 26 | 3 | 26.71875 | 32 |
| VBC | F | 40 | 3 | 6.0625 | 31 |
| VBC | F | 47 | 3 | 22.71875 | 30 |
| VB | M | 12 | 4 | 75.875 | 32 |
| VB | M | 26 | 4 | 88.5 | 32 |
| VB | M | 40 | 4 | 71.15625 | 29 |
| VB | M | 47 | 4 | 80.3125 | 26 |
| VB | F | 12 | 4 | 93.15625 | 32 |
| VB | F | 26 | 4 | 72.9375 | 32 |
| VB | F | 40 | 4 | 51.4375 | 32 |
| VB | F | 47 | 4 | 41.59375 | 31 |
| VBC | M | 12 | 4 | 18 | 32 |
| VBC | M | 26 | 4 | 29.53125 | 32 |
| VBC | M | 40 | 4 | 16.78125 | 28 |
| VBC | M | 47 | 4 | 28.9375 | 21 |
| VBC | F | 12 | 4 | 24.65625 | 32 |
| VBC | F | 26 | 4 | 20.875 | 32 |
| VBC | F | 40 | 4 | 15.6875 | 31 |
| VBC | F | 47 | 4 | 19.5625 | 28 |

*Assay began with 32 flies for each treatment. The sample sizes reported above reflect only the data from flies that survived and were analysed.

**Table S2. Descriptive statistics of locomotor activity for VB and VBC populations at all age points in Experiment 2**

| Day | **VB** | | | **VBC** | | |
| --- | --- | --- | --- | --- | --- | --- |
|  | Sample size | Average Activity (counts/h) | 95% CI | Sample size | Average Activity (counts/h) | 95% CI |
| **12** | 54 | 187.6 | 14.4 | 57 | 50.7 | 6.8 |
| **26** | 53 | 147.3 | 15.8 | 54 | 40.8 | 6.7 |
| **33** | 50 | 144.7 | 18.3 | 54 | 34.3 | 6.1 |
| **40** | 44 | 111.5 | 18.4 | 51 | 30.9 | 6.0 |
| **47** | 27 | 94.9 | 19.5 | 36 | 25.3 | 5.8 |

**Table S3. Statistics regarding lifespan measurements of VB and VBC population**

| Measurement variable | VB | | VBC | | ANOVA *F* value | ANOVA *p* value |
| --- | --- | --- | --- | --- | --- | --- |
|  | Average | 95% CI | Average | 95% CI |  |  |
| Lifespan  (days) | 45.7 | 2.29 | 50.4 | 3.18 | 4.517 | 0.036 |

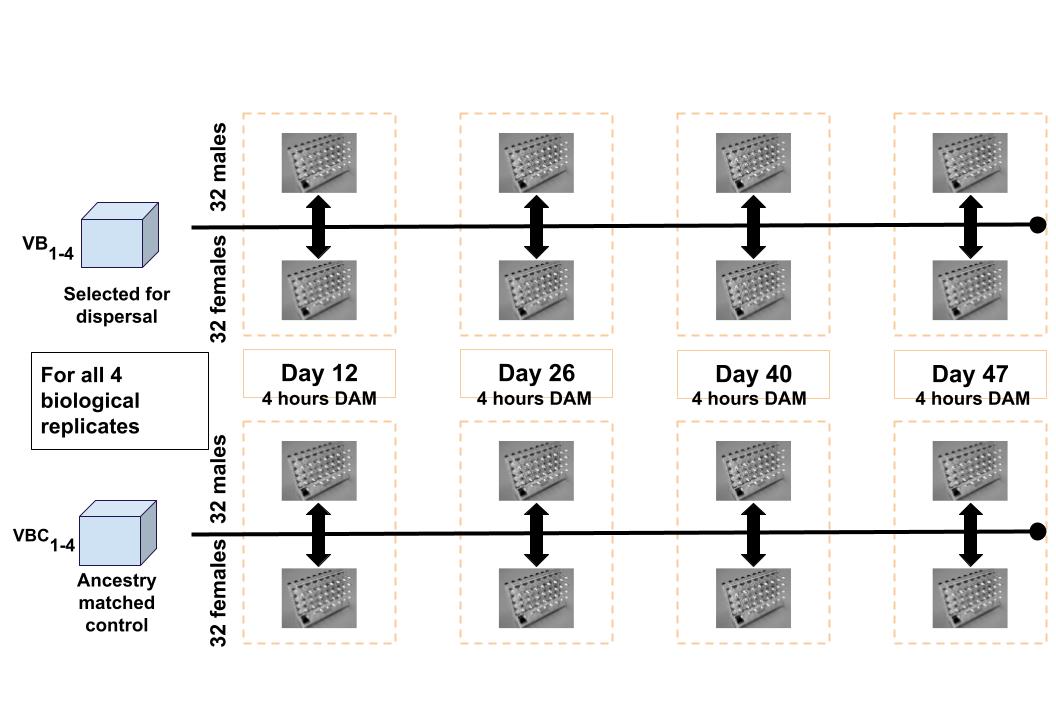

**Figure S1.** Schematic representation of assay design for Experiment#1 i.e., the cross-sectional locomotor activity assay where locomotor activity of a set of individuals from the population was measured at the give age points.

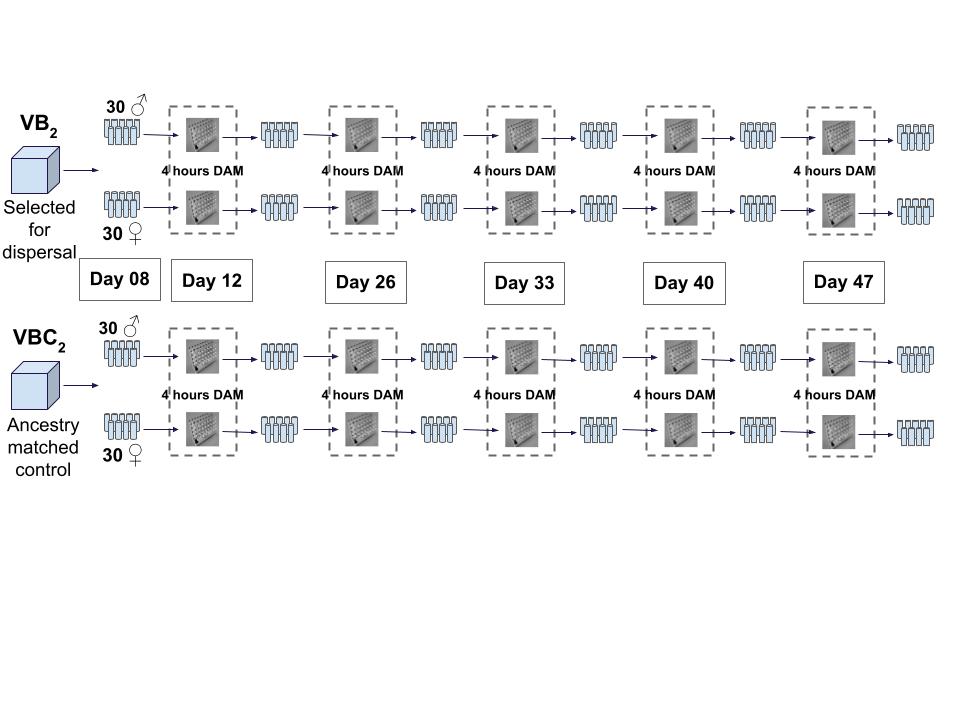
**Figure S2.** Schematic representation of assay design for Experiment#2 i.e., the longitudinal locomotor activity assay where same individuals were followed throughout the experiment. Note that this experiment was conducted only in one of the dispersal selected populations along with its ancestry matched control population.

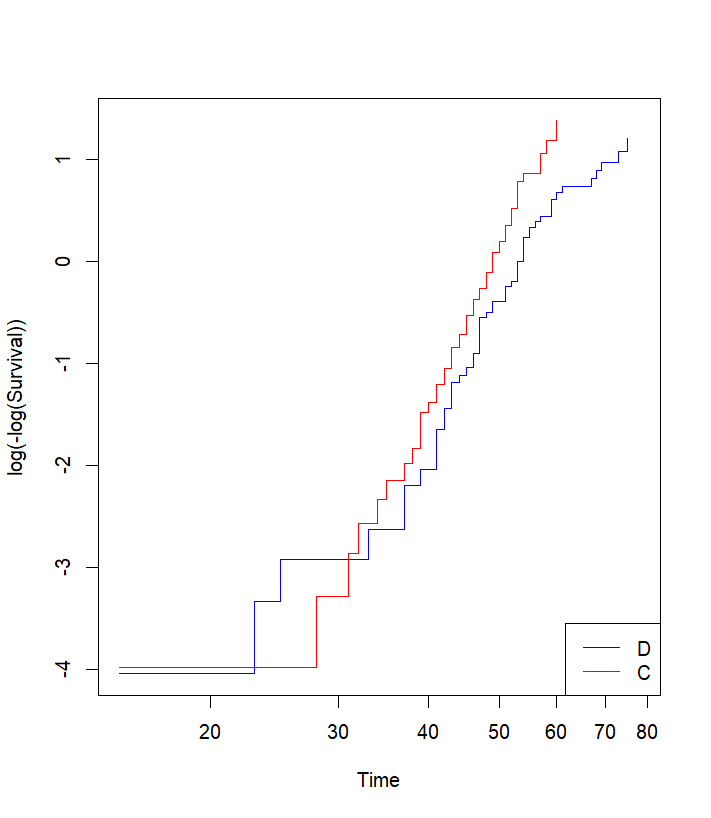

**Figure S3. Log-minus-log survival plot for D and C populations.** The log-minus-log survival plot shows the relationship between survival time and the cumulative hazard for the dispersal selected D (blue line) and control C (red line) populations. The parallelism of these lines is used to assess the proportional hazards assumption in Cox proportional hazards model, with the relatively parallel lines suggesting that the proportional hazards assumption is reasonably met.

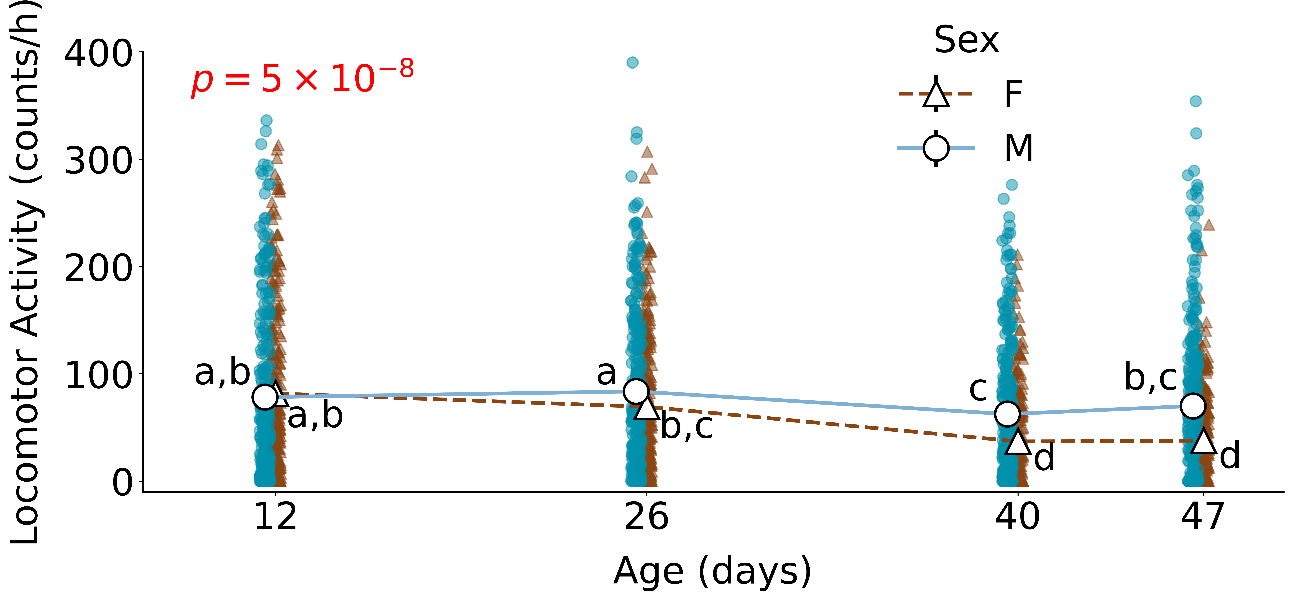

**Figure S4.** Locomotor activity of male and female flies across age in the cross-sectional experiment. Average locomotor activity of females in later age points is significantly less than males. *p* value corresponding to Sex × Age interaction is written in red font. The teal and brown scatter points represent the data for all the replicates of male and female flies respectively. Different lower-case alphabets denote statistically significant differences (Tukey *p* <0.05).

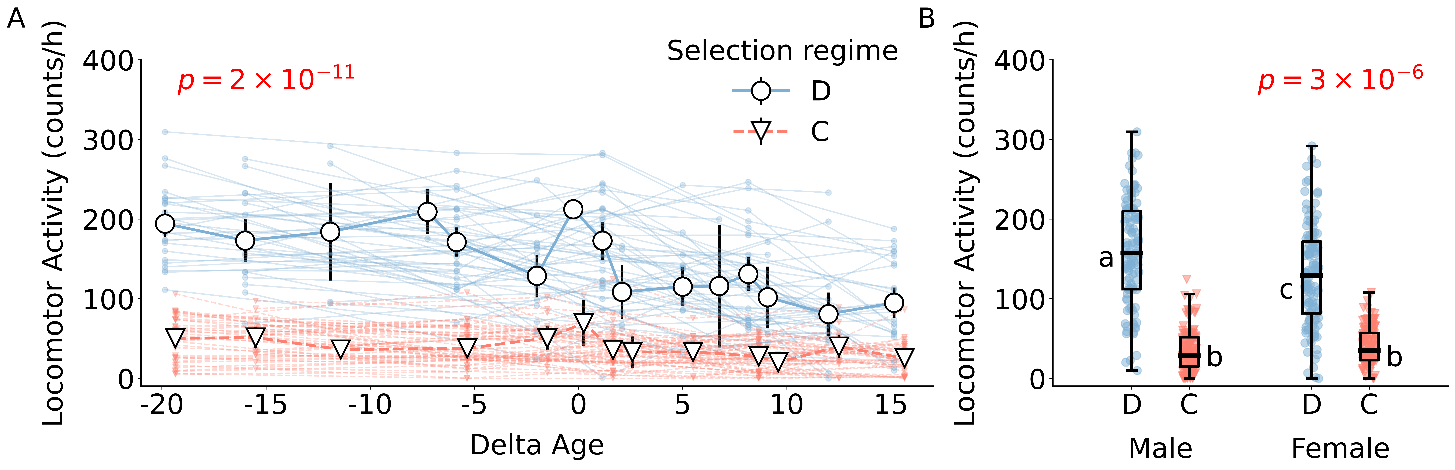

**Figure S5. Age-and sex-dependent locomotor activity profile from longitudinal data.**
(A) Mean (± 95% CI) locomotor activity of individuals from dispersal selected population D (circle) and its ancestry-matched control population C (triangle) are plotted against delta age. Activity level of each individual at the tested age-point of dispersal selected and control populations are represented as blue and red scatter points respectively. Scatter points for same individual are connected by solid line. *p* value for Selection × Delta age interaction is mentioned in red font. (B) Box-plot representing locomotor activity of individuals from dispersal-selected D and its ancestry-matched control C populations across males and females. The blue and red scatter points represent the data for all the replicates of D and C populations respectively. The edges of the box denote 25^th^ and 75^th^ percentiles, while the black solid line represents the median. *p* value for Selection × Sex interaction is mentioned in red font. Different lower-case alphabets denote statistically significant differences (p<0.05). Some of the error bars are not visible due to their small size.

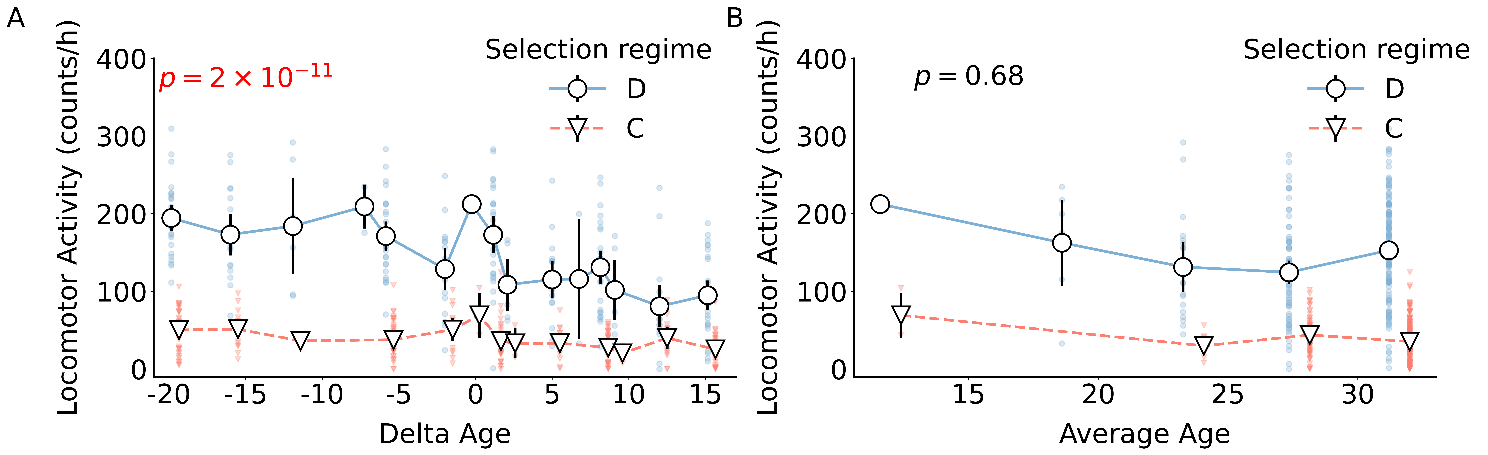

**Figure S6.** Mean (± 95% CI) locomotor activity of individuals from dispersal selected population D (circle) and its ancestry-matched control population C (triangle) are plotted against Average age. Activity level of each individual at the tested age-point of dispersal selected and control populations are represented as blue and red scatter points respectively. Scatter points for same individual are connected by solid line. *p* value for Selection × Average age interaction is mentioned in black font.
